# The influence of data type and functional traits on native bee phenology metrics: Opportunistic versus inventory records

**DOI:** 10.1101/2020.04.16.044750

**Authors:** Joan M. Meiners, Michael C. Orr, Riemer Kristina, Griswold Terry, Juniper L. Simonis

**Author notes:** First three authors should be considered joint first authors (Joan M. Meiners, Michael C. Orr, Riemer Kristina).

## Abstract

Efforts to understand activity patterns of bees, our most important pollinators, often rely on opportunistically collected museum records to model temporal shifts or declines. This type of data, however, may not be suitable for this purpose given high spatiotemporal variability of native bee activity. By comparing phenological metrics calculated from intensive systematic inventory data with those from opportunistic museum records for bee species spanning a range of functional traits, we explored biases and limitations of data types to determine best practices for bee monitoring and assessment. We compiled half a million records of wild bee occurrence from opportunistic museum collections and six systematic inventory efforts, focusing analyses on 45 well-represented species that spanned five functional traits: sociality, nesting habits, floral specialization, voltinism, and body size. We then used permutation tests to evaluate differences between data types in estimating three phenology metrics: flight duration, number of annual abundance peaks, and date of the highest peak. We used GLMs to test for patterns of data type significance across traits. All 45 species differed significantly in the value of at least one phenology metric depending on the data type used. The date of the highest abundance peak differed for 40 species, flight duration for 34 species, and the number of peaks for 15 species. The number of peaks was more likely to differ between data types for larger bees, and flight duration was more likely to differ for larger bees and specialist bees. Our results reveal a strong influence of data type on phenology metrics that necessitates consideration of data source when evaluating changes in phenological activity, possibly applicable to many taxa. Accurately assessing phenological change may require expanding wild bee monitoring and data sharing.

## Introduction

Accurately estimating species phenology is central to understanding ecological systems (J. Forrest & Miller-Rushing, 2010; Inouye, 2008; Nakazawa & Doi, 2012). When species are active determines the abiotic conditions they face; the identity, quality, and quantity of resources available to them; and the specific competitors and predators they encounter. Yet the relative timing of ecosystem components can be difficult and costly to assess. The activity pattern of any given species may vary both temporally (across years) and spatially, in response to a cacophony of abiotic and biotic conditions, which themselves fluctuate at various scales (de Keyzer et al., 2017). The more environmentally-sensitive and species-rich the taxon of interest, the more complicated it can be to determine the magnitude, or even the direction, of any generalized phenological trends (de Keyzer et al., 2017; Primack et al., 2009). On top of these biological considerations, the amount and type of data necessary to evaluate phenological patterns or navigate known biases remains unclear (de Keyzer et al., 2017; Isaac & Pocock, 2015; Miller-Rushing et al., 2010; Strien et al., 2008). A better understanding of these consequential uncertainties is necessary to reliably determine the effects of environmental change on both natural and managed systems.

Native bees include highly variable and diverse taxa that are of particular interest for phenological studies due to their value as pollinators and their vulnerability to ecosystem change (Fabina et al., 2010; Ogilvie & Forrest, 2017; Rafferty & Ives, 2011). Many native bees are solitary, and respond to a host of environmental cues to time emergence from overwintering nests, including changes in soil moisture from snowmelt or precipitation, and temperature, which can influence the rate at which larvae exhaust food supplies and undergo metamorphosis (Danforth, 1999; Helm et al., 2017; Michener, 2007). The interplay of species-specific demography and responses to abiotic emergence cues is poorly understood for the majority of native bee species, and may or may not align with factors determining local seed germination and bloom time (Aldridge et al., 2011; Danforth, 1999; Miller-Rushing et al., 2010). The potential for temporal mismatches between pollinators and their host plants, which could result in inadequate pollination for plant reproduction and nutritional deficits for bees, has inspired many recent studies of phenological shifts. Studies of this phenomenon have returned mixed results, with evidence both of problematic disruptions of historical pollinator-plant relationships (Aldridge et al., 2011; Burkle et al., 2013; Robbirt et al., 2014; Schenk et al., 2018) and of inconclusive or parallel shifts in emergence and bloom time (Bartomeus et al., 2011; J. R. K. Forrest, 2015; J. R. K. Forrest & Thomson, 2011; Ogilvie & Forrest, 2017). Little attention, however, has been paid to how the type, quantity, or quality of data used to measure phenology in bees – or other species – may produce conflicting conclusions.

To assess the vulnerability of plant-pollinator relationships to climate change, we first need to examine whether data used to assess bee phenology accurately reflects real changes in bee activity, as opposed to noise from unevenly-sampled biological variability, biased collecting protocols, or sample-size limitations of the data. Sampling diverse organisms at large spatial and temporal scales can be an incredibly laborious and expensive process. As a result, our knowledge of bee trends necessarily draws from patchy and inconsistently-collected data (Meiners et al., 2019). Understanding what available data can and cannot tell us about phenological trends over time is an oft-overlooked prerequisite for accurately modeling the status, trends, and impacts of wild bee abundance, as well as forecasting declines, range shifts, risks to network stability, and vulnerability of particular species to climate change (Biesmeijer et al., 2006; Forrest, 2015; Meiners et al., 2017; Potts et al., 2016). This oversight is apparent in pollinator studies but may impact phenological analyses of other taxa as well. Furthermore, testing and communicating all sources of error and uncertainty is an important step in advancing scientific understanding and maintaining public trust and investment in the ability of science to measure and mitigate changes in our natural world.

Data available to researchers interested in large-scale animal activity trends can generally be divided into two types: “opportunistic” and “inventory”. “Opportunistic” data usually consist of records compiled from museum specimens belonging to specific groups of interest that are the result of various disparate project collections, often of unknown and unspecified purposes. With millions of records served from curated, digitized museums to public, online hubs, opportunistic data are a rich resource of natural history information uniquely voluminous and useful for a range of research questions (Lister & Climate Change Research Group, 2011). They are also known, however, to contain biases and limitations derived from their unstandardized and composite origins (Isaac & Pocock, 2015), and to lack metadata that would allow for easy exclusion of biased records from small projects with specific objectives, such as sampling only the bees visiting a particular plant. “Inventory” data, on the other hand, are collected in a systematic manner, often for the explicit purpose of answering broad questions about place-specific biodiversity patterns and community processes. A standardized protocol for conducting inventories of native bee communities was established in 2003 by a group of melittologists (LeBuhn et al., 2003), and has been used in many inventories of bees in natural areas (Griswold et al., 1999; Kuhlman & Burrows, 2017; Meiners, 2016; Messinger, 2006). The expense and effort required to follow a systematic inventory protocol is higher, but the assumption is that inventory records have fewer biases, resulting in superior estimates of bee activity, floral reliability (Wright et al., 2015), and baseline community patterns against which evaluations of future change can be measured (Meiners et al., 2019). Despite these core data type differences, however, both opportunistic and inventory data have been used interchangeably in studies of native bee phenology without an assessment of their relative suitability for the task. Drawing mistaken conclusions from data that is flawed, incomplete, or was collected for another purpose may result in mismanagement of natural resources, misdirected sampling efforts, and missed opportunities to harness the full power of both opportunistic and inventory datasets.

We use data from six systematic bee inventories and approximately a quarter million opportunistic museum records collected over twenty years to compare estimates per data type of three phenology metrics for forty-five abundant native bee species. To assess the possibility of extrapolating conclusions to additional species, we also examine trends related to five functional life history traits, which recent research has shown to be predictive of native bee rates of decline (Bartomeus et al., 2013), vulnerability to insecticides (Brittain & Potts, 2011), response to anthropogenic disturbance (Williams et al., 2010), and pollinating behavior (Pisanty & Mandelik, 2015). With this approach, we seek to answer two central questions: 1) can opportunistic data produce parameter estimates of native bee species phenology that are statistically equivalent to more expensive inventory data?, and 2) if phenology metrics differ between data types, are there patterns associated with functional traits that could be useful indicators of which species are more susceptible to erroneous phenology estimates? In answering these questions, we seek to improve the utility of natural history collection data, the determination of native bee trends and conservation practices, and the broad reliability of phenology results.

## Methods

### Data, species, and trait selection

Opportunistic and inventory data were sourced from the USDA-ARS National Pollinating Insects Collection (NPIC) (USDA-ARS, 2016). We defined opportunistic data as specimen records which were collected irrespective of each other or standard protocols, instead typically irregularly targeting regions, specific floral resources, or bee taxa of special interest. We defined inventory data as records arising from standardized efforts to systematically document bee species richness in specific geographic areas across the local season(s) of bee activity. Six studies with collections housed at the NPIC fit this definition of inventory data. These were systematic inventory studies located at: Carlsbad Caverns National Park (*Griswold & Ikerd unpublished data*); Clark County, Nevada (Griswold et al., 1999); Grand Staircase-Escalante National Monument (Carril et al., 2018; Messinger, 2006); Pinnacles National Park (Meiners, 2016; Meiners et al., 2019); and Yosemite National Park (*Griswold & Ikerd unpublished data*). All six inventory studies were conducted between 1996 and 2012 following protocols similar to that outlined in LeBuhn et al. (2003), and shared at dx.doi.org/10.17504/protocols.io.wfhfbj6 from Meiners et al. (2019).

We restricted the temporal and spatial range of our study within reason, while keeping our dataset large to limit the phenological variability introduced solely by spatiotemporal factors. For both inventory and opportunistic data, we only used specimen records that met all of the following criteria: 1) identified to a valid species, 2) collected between 1990 and 2015 in the USA or Canada, and 3) contained complete and reliable georeferencing. Data cleaning to meet these criteria was conducted in R (R Core Team, 2015). To ensure sufficient sample sizes for species-level comparisons between data types, we excluded any species with fewer than 180 occurrences for each data type, retaining only the fifty most abundant species shared between the two data sets.

Once we finalized our list of fifty species, we conducted literature searches and expert surveys to assign them into categories of five pre-selected life history traits that literature searches and expert consensus suggested have relevance to phenological trends: *body size, sociality, floral specialization, nest location*,and *voltinism* (Araújo et al., 2004; Heithaus, 1979; Osorio-Canadas et al., 2016; Rodriguez-Girones & Bosch, 2012; Williams et al., 2010). We used a Keyence digital microscope to measure *body size*as the average inter-tegular distance (between wing bases) for five female specimens of each species, following the method specified by Cane (1987). Based on species-specific literature searches, we categorized the *sociality* of a bee species as either 1) solitary, or 2) social, which included bee species that can be described as eusocial, communal, and primitively social, or 3) unknown (our list of fifty did not include any cleptoparasitic species). We noted whether a species was considered a *floral specialist* in the literature by a simple 1) yes, 2) no, or 3) unknown. We noted *nest location*as a binary trait, with species categorized as nesting primarily 1) above ground or 2) below ground. Due to a lack of published information, we classified *voltinism* based on a survey of expert opinion into the following classes: 1) univoltine (one generation), 2) multivoltine (>1 generation), 3) social (since these species replace members throughout the season but not in the same way as multiple generations of solitary bees in a single season), and 4) unknown.

### Final Dataset Specimen, Species, and Trait Composition

The final dataset contained 104,101 bee occurrence records, of which 71,152 were from inventory collections and 32,949 were opportunistically collected. From the original fifty species, we removed five from the dataset because they are either: 1) commonly managed (*Apis mellifera)*; 2) have an unusual, socially-parasitic life history (*Bombus insularis*); or 3) could not reliably be distinguished in females (*Agapostemon angelicus, Agapostemon texanus, Agapostemon angelicus/texanus*), resulting in a final set of 45 species (Table 1).

**Table 1.**
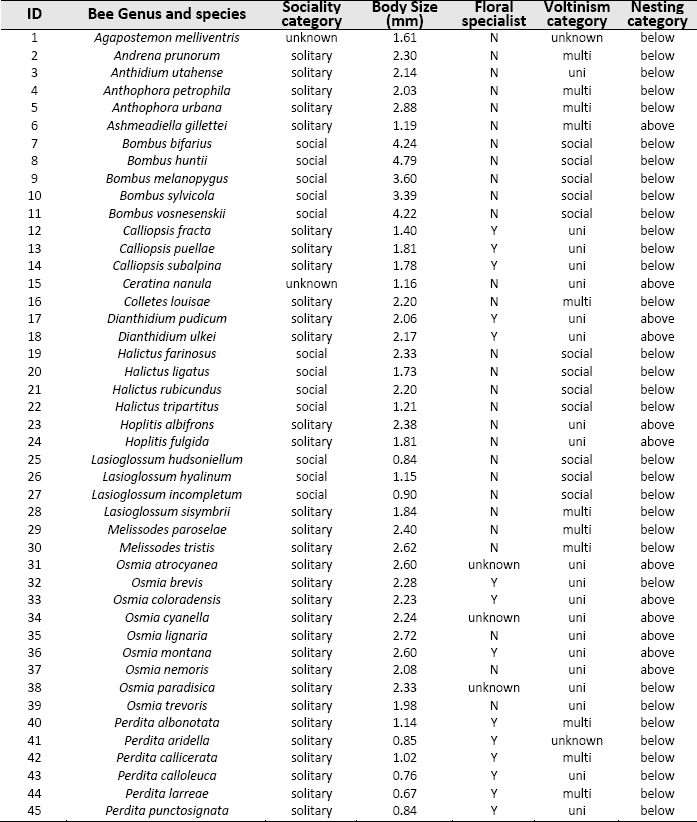
Species (N=45) selected for analysis based on number of records (N >180 for each data type) and spread of representative traits (N = 5). Sources of trait information are included in SI.

The final 45 species selected for phenology metric analyses showed a relatively even spread of traits. Some trait category assignments for certain species were impossible to assign based on current knowledge and remain labeled as “unknown” in our trait dataset (Table 1). All assigned trait categories were represented by at least 12 out of 45 total species.

### Calculation of phenology metrics

We identified three measurable metrics of bee phenology that would be useful and reliable for quantitatively estimating changes in patterns of bee species activity over time: 1) flight duration, or the number of days in a year the bee species was active; 2) clusters, or the number of distinct peaks in abundance in a year; and 3) the date of a bee species’ highest annual peak in abundance (Fig. 1). We defined flight duration as the middle 90% of occurrences, removing the upper and lower 5% of values to eliminate outliers that may represent unusual activity in any given year. We determined the number of clusters in a set of occurrences, with a maximum possible of three clusters, using a gap statistic. We then used kmeans clustering to find the location along the day-of-year axis of all clusters. The cluster with the highest value on the density plot was chosen as the date for the greatest abundance of occurrences. We calculated these three metrics twice for each species, once each for all occurrences from the inventory data type and once for all the opportunistic data type occurrences. In order to have a single number for each metric that showed how different they were for the two data types, we calculated a test statistic for each of the three metrics. We chose the test statistic as the absolute difference between the opportunistic and inventory metric, so that each species had three test statistics, one each for flight duration, number of clusters, and location of greatest abundance.

**Fig. 1.**
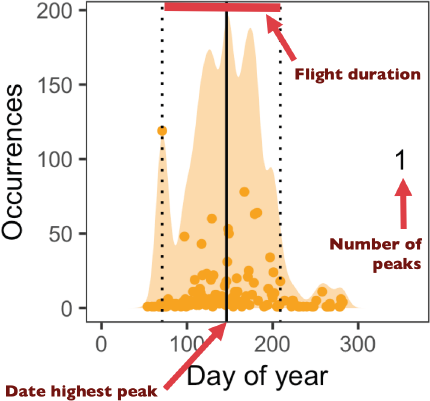
Visualization of three phenology metrics calculated for each of 45 species and two data types. Number of occurrences each day of the year are plotted as orange points, which are also represented as a density plot by the orange smoothed histogram. Vertical black dotted lines indicate the beginning and end of flight duration, the number shows the number of clusters in occurrences, and the location of the maximum peak in occurrences is represented by a vertical solid black line. In subsequent graphs, a gray line indicates secondary peaks in abundance.

**Fig. 2.**
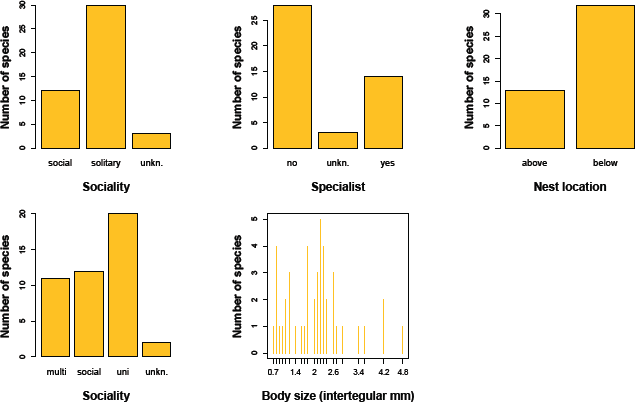
Distribution of 45 species across categories and values of five functional traits.

In order to determine if these test statistics indicated that there was a substantial difference between phenology patterns for data types, we compared these observed test statistics to a set of simulated test statistics that came from shuffling the data. We randomly shuffled the data type labels for all occurrences of each species, retaining the same relative number of opportunistic and inventory labels for each species. We then recalculated the three phenology metrics for the two data types and the test statistic, so that each species had three simulated test statistics. Finally, we repeated this process 1000 times, so that each species had a distribution of simulated test statistics.

To determine if the observed test statistics were statistically significantly different than the distribution of simulated test statistics, which would indicate that data type mattered for that phenological pattern, we calculated a p-value based on the number of simulated test statistics that were greater than the observed one. We used an alpha cut-off of 0.05, and each species had one p-value for each of the three phenology metrics. Given the multitude of pairwise comparisons, we also include a more stringent alpha cut-off of 0.001, which is the lowest value that can be achieved given the number of permutations.

### Modeling influence of functional traits

After assigning a category value to each bee species for each of the five selected functional traits, as described above, we used generalized linear models to assess the influence of functional traits on significant differences between data types from permutation tests for each of the three identified bee phenology metrics (Fig. 1). This evaluation of the influence of functional traits on data type significance was only conducted for species with complete trait category information (Table 1). Species with “unknowns” were removed, and voltinism levels “social” and “multi” were ultimately combined so that all categorical traits were binary variables. All data manipulation, plotting, and statistical tests were conducted in the R statistical package (R Core Team, 2015).

## Results

### Phenology Metrics by Data Type

With three phenology metrics for each of 45 species, we compared a total of 135 pairs of phenology variable calculations based on data type. We found significantly different values depending on which data type was used (inventory or opportunistic) in 87 out of 135 cases, which represents 64% of the possible total, much higher than the 5% expected under a null hypothesis and assuming a 5% alpha. The date of highest peak in abundance was the metric with the greatest number of value discrepancies due to data type: the date of the seasonal peak was significantly different depending on which dataset was used for 40 out of 45 species (89%, Fig. 3). Flight duration, or the number of days a species was active, differed based on data type for 34 out of 45 species (76%, Fig. 3). And the number of clusters, or distinct peaks in abundance, was different between data types for 15 out of 45 species (33%, Fig. 3). It should be noted, however, that the number of clusters was the least reliable of the three phenology metrics, due to limitations of the gap statistic used to calculate it, sensitivity to variable collection efforts over time in the opportunistic data, and the narrow range of options between just one and three for number of clusters detected. We considered other options for calculating number of clusters but found the gap statistic to be the most defensible, if still flawed.

**Fig. 3.**
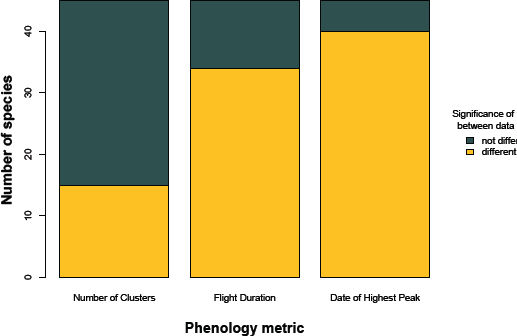
Number of species, out of 45, for which values of each of three phenology metrics differed significantly (*alpha* =0.05) based on the data type used to calculate them.

All 45 species had at least one metric that was significantly different depending on data type (Table 2). Ten out of 45 species had significantly different results for all three phenology metrics depending on the type of data used to evaluate them. Occurrence curves for each species and data type illustrate the comparison between inventory and opportunistic data types of the three phenology metrics (Fig. 4). The species *Ceratina nanula*, for example, differed in flight duration between data types, seen as the width of the x-axis between dotted lines, but had statistically similar results for the date of the highest peak and the number of clusters in abundance (Fig. 4, top left). *Lasioglossum sisymbrii* had the same number of clusters in both inventory and opportunistic datasets, but different values for both flight duration and date of the highest peak, shown by the solid vertical line on the plot (Fig. 4, top right). Two species of *Osmia* had different results for the number of clusters reported by the gap statistic (Fig. 4, middle row), as well as either a different flight duration or different date of highest peak depending on data type. As mentioned above, ten species differed in all three metrics between data types, as illustrated by *Lasioglossum hudsoniellum* and *Anthophora urbana* (Fig. 4, bottom row). Because significance of the difference between data type was based on 1000 different permutations of the occurrence records, not all figures showing species-level results match the reported conclusions.

**Fig. 4.**
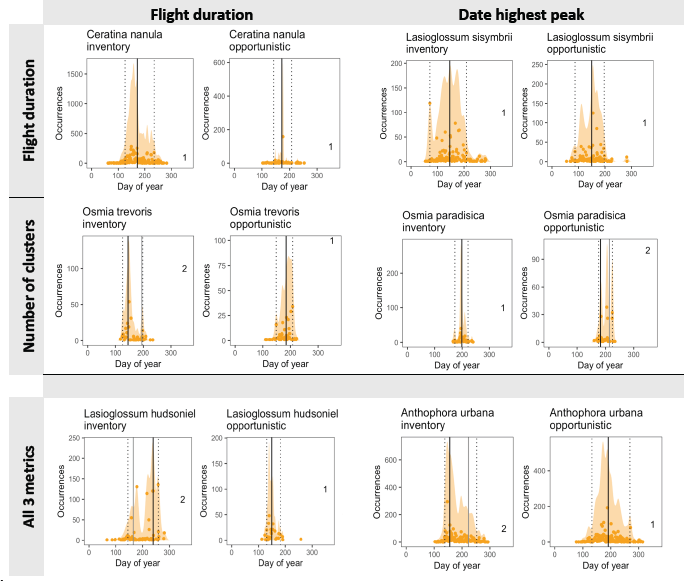
Examples of permutation tests results comparing phenology metrics of bee species (N=45) occurrence distributions between two data types. In each cell panel, inventory data is plotted on the left and opportunistic for the same species is plotted data on the right. Column and row names indicate phenology metrics that were found to be significantly different between data types for the species plotted in those cells. In the top left, only flight duration was different between data types, while in the top right both flight duration and date of the highest peak were different, and all three metrics were different for both species in the bottom row. Occurrence curves showing results for all 45 species are in Supporting Information.

**Table 2.** Incidences where each metric returned significantly different results between two data types for each of 45 species (* indicates *p*<0.05 and ** indicates *p*<0.001).

| ID | Bee Genus and species | Number of clusters | Flight duration | Date highest peak | # metrics differing |
| --- | --- | --- | --- | --- | --- |
| 1 | *Agapostemon melliventris* |  | ** | * | 2 |
| 2 | *Andrena prunorum* |  | ** | ** | 2 |
| 3 | *Anthidium utahense* |  | ** | ** | 1 |
| 4 | *Anthophora petrophila* |  |  | * | 1 |
| 5 | *Anthophora urbana* | ** | ** | ** | 3 |
| 6 | *Ashmeadiella gillettei* |  | ** | ** | 2 |
| 7 | *Bombus bifarius* |  | ** | ** | 2 |
| 8 | *Bombus huntii* |  |  | * | 1 |
| 9 | *Bombus melanopygus* |  |  | ** | 1 |
| 10 | *Bombus sylvicola* | ** |  | ** | 2 |
| 11 | *Bombus vosnesenskii* |  | ** | ** | 2 |
| 12 | *Calliopsis fracta* | ** | ** |  | 2 |
| 13 | *Calliopsis puellae* |  |  | * | 1 |
| 14 | *Calliopsis subalpina* |  | ** | * | 2 |
| 15 | *Ceratina nanula* |  | ** |  | 1 |
| 16 | *Colletes louisae* |  | ** | ** | 2 |
| 17 | *Dianthidium pudicum* | * | ** | * | 3 |
| 18 | *Dianthidium ulkei* |  | ** | ** | 2 |
| 19 | *Halictus farinosus* |  | ** | ** | 2 |
| 20 | *Halictus ligatus* |  | ** | ** | 2 |
| 21 | *Halictus rubicundus* |  |  | * | 1 |
| 22 | *Halictus tripartitus* | ** | ** | ** | 3 |
| 23 | *Hoplitis albifrons* |  | ** | ** | 2 |
| 24 | *Hoplitis fulgida* |  | * | ** | 2 |
| 25 | *Lasioglossum hudsoniellum* | ** | ** | ** | 3 |
| 26 | *Lasioglossum hyalinum* | * | ** | ** | 3 |
| 27 | *Lasioglossum incompletum* | ** | * | ** | 3 |
| 28 | *Lasioglossum sisymbrii* |  | ** | * | 2 |
| 29 | *Melissodes paroselae* |  | ** | ** | 2 |
| 30 | *Melissodes tristis* |  | * |  | 1 |
| 31 | *Osmia atrocyanea* | ** | ** | ** | 3 |
| 32 | *Osmia brevis* |  |  | ** | 1 |
| 33 | *Osmia coloradensis* |  | ** | ** | 2 |
| 34 | *Osmia cyanella* |  |  | ** | 1 |
| 35 | *Osmia lignaria* |  | * | ** | 2 |
| 36 | *Osmia montana* |  |  | ** | 1 |
| 37 | *Osmia nemoris* |  | ** | ** | 2 |
| 38 | *Osmia paradisica* | ** |  | ** | 2 |
| 39 | *Osmia trevoris* | * | ** |  | 2 |
| 40 | *Perdita albonotata* | * | * | * | 3 |
| 41 | *Perdita aridella* | ** | ** | ** | 3 |
| 42 | *Perdita callicerata* |  |  | ** | 1 |
| 43 | *Perdita calloleuca* |  | * | ** | 2 |
| 44 | *Perdita larreae* |  | ** |  | 1 |
| 45 | *Perdita punctosignata* | * | ** | ** | 3 |
| **Totals** |  | **15** | **34** | **40** | **87** |
| **Percent of possible** |  | **33** | **76** | **89** | **64** |

### Relationship of Functional Traits to Data Type Significance

For the group of forty species without any unknown trait values, body size was a significant (*p*= 0.047) predictor of whether flight duration would differ between data types, with larger bees being more likely to have different results for seasonal activity length depending on which dataset was used (Table 3). Generalized linear model results also found body size to be a marginally significant (*p* = 0.055) predictor of whether the number of clusters would differ between data types, with larger bees more likely to return different number of clusters depending on whether opportunistic or inventory records were used to calculate them. Floral specialization was also a marginally significant (*p* = 0.051) predictor of difference between data type in flight duration, with species designated as floral specialists more likely to return different values for number of days they were active over a season depending on data type (Table 3). The likelihood for date of the highest peak to differ between opportunistic and inventory data was not significantly related to any of the five functional traits.

**Table 3.**
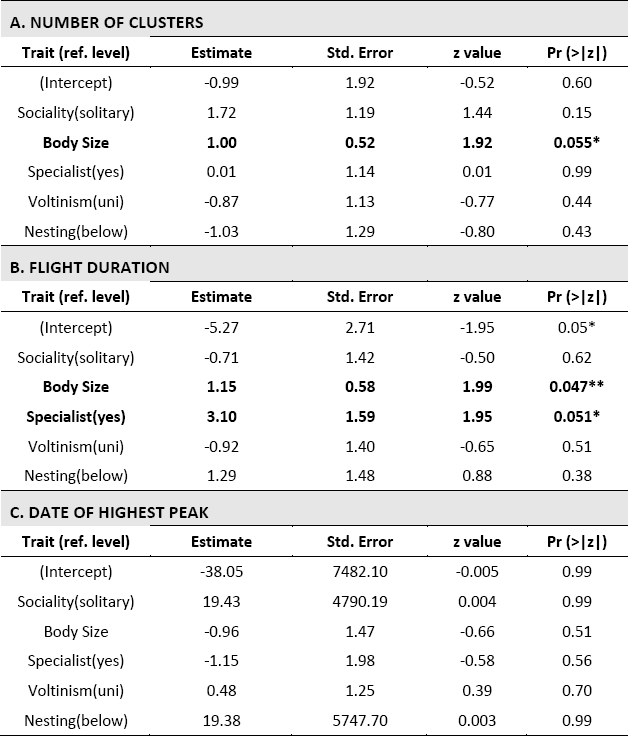
Results of generalized linear models evaluating significance of species (N=40, those without any unknown traits) functional traits on the difference between data types in calculating values of three phenology metrics.

## Discussion

We found widespread and significant differences in estimates of phenology metrics depending on the type of data used to calculate them. Out of 45 tested native bee species spanning a range of life history traits, one hundred percent had a significantly different value for at least one key phenology metric depending only on the type of data used to calculate it. This result should raise concerns about the influence of data source on our understanding of patterns and changes in phenology, not only for pollinators but potentially for other taxa studied using compiled museum records. With the high natural variability of many small organisms already obscuring measurable signals of behavior, adding noise to phenology models by using messy or inappropriate data may confound phenological estimations to the point that they become uninformative. If biases are consistent and directional, phenology studies that do not take into account the influence of data may even report patterns opposite the truth.

Our study reveals an urgent need to ensure that the data used for evaluating changes in phenology, not only of native bees but likely of many other organisms as well, are of sufficient quality to produce reliable results. Comparing metrics over time that are not compatible – for example, checking for changes in the date of peak species activity by comparing recent inventory records to older opportunistic records from historical study sites – may add noise instead of clarity to collective efforts to detect real changes in phenology, or may create false impressions of a pattern. Critical interaction mismatches between any organisms reliant on each other could be obscured. Such misleading results may also hinder scientific progress and conservation efforts, erode public trust in science, and dilute the gravity of warnings about pollinator declines and other environmental changes.

The implications of our study may be relevant in many systems but are certainly of consequence as applied to native bees. Since plant reproductive success depends on the timing of local pollinator activity, recently found to be shifting with climate change (Aldridge et al., 2011; Burkle et al., 2013; Robbirt et al., 2014; Schenk et al., 2018), phenology research on biodiverse networks of coexisting native bee species and their hosts is highly complex but vitally important. Even with the best data, results from one study may provide limited insight for patterns in another area, species, or time. Primack et al. (2009), for example, conducted phenology studies for twelve taxa using long-term, spatially-expansive, and systematically-collected data, and still found results to be highly variable and hard to interpret due to species-level variability. With the explosion of technology, museum data-basing efforts, and the open science movement, the availability of data to ask important questions about bee species phenology is entering a new frontier. It is, therefore, necessary to also update our understanding of data and methodological limitations before extrapolating findings across regions or species, potentially muddying conclusions about the high-profile issues of bee decline and worldwide loss of pollination services (Biesmeijer et al., 2006; Burkle et al., 2013; Goulson et al., 2015).

Despite the overall increased availability of data, records are still very limited for many taxa, and that is where common parameters like functional traits can be useful. Until further technological advances in bee species identification and specimen processing make it feasible to obtain sufficient data to evaluate phenological trends for a majority of cryptic or rare native bee species, efforts to identify unifying variables that correspond to data type reliability or phenological variability will be relevant. Our result that the importance of having high-quality data increases with increasing body size and increasing floral specialization for native bees, for instance, can help guide studies of smaller groups of species when deciding how to allocate data collection resources. Larger bees can emerge earlier in the season than smaller-bodied bees, due to their greater ability to generate and maintain elevated body temperatures under cold conditions (Osorio-Canadas et al., 2016). Being active earlier in spring may make spring-flying larger bees more variable in the interannual timing of their activity as the date of snowmelt and first bloom also vary. Likewise, being tied to a particular floral group as a foraging specialist bee species may require greater flexibility in emergence time and a stronger reliance on specific environmental cues to time emergence that may make specialist phenology more difficult to evaluate. Opportunistic data yielded inconsistent values for flight duration for larger-bodied and specialist bees in our study. Knowing, as a result, that inventory data is more appropriate for these species allows for cleaner interpretation of their behavior. Likewise, it is useful to know from this result that opportunistic data may be more appropriate for estimating phenology metrics for smaller species and floral generalists, at least where many records are available. In these ways, our functional trait model illustrates how exploring limitations of data types can have both biological and statistical value.

Our study does not seek to undermine the great importance or value of natural history or museum collections, but rather to explore and illuminate best practices for data use in the study of phenology. The appropriate source of data may depend entirely on the nature or scale of the question being asked, or the level of specificity desired. Systematic inventories of native bee fauna provide ideal data for understanding bee ecology, but are hugely expensive and time-consuming, and should not take the place of opportunistic data for every research endeavor. In some cases, such as when gauging phenological changes across decades, it may not be possible to rely on inventory data, but the limitations of the data available must still be understood, because the best available data may fail to provide the correct answers, regardless of the methods employed. There is much to be gained from appropriate use of opportunistic data to estimate metrics of species phenology, and much to be lost from ignoring it. The influence of data type on phenology estimation is likely important for many other taxa with spotty records and high inherent variability. Incorporating measures of data bias and associated relevance of functional traits to guide interpretation of results may benefit the study of phenology and ecology in a myriad of ways.

While we improve our use of data, we must also continue expanding our knowledge base. Natural history collections across the world are struggling to attain the financial, institutional, and cultural support required to develop, curate, document, and digitize museum collections. Improving the flow of high-quality data records from diverse areas and time periods is an important step in alleviating data bias and improving our understanding of phenology. Expanding and further standardizing inventory efforts will also be important. The majority of broad-scale bee phenology studies have taken place in montane or agricultural landscapes, or on social, cavity-nesting bees, leaving other environments and guilds poorly understood (Bosch & Kemp, 2002; CaraDonna et al., 2014; Hanley et al., 2015; Klein et al., 2007; Ogilvie & Thomson, 2015; Winfree et al., 2011). Since we know that the extrapolation of conclusions about phenology patterns across space, species, and data type is flawed (Davis et al., 2010; Primack et al., 2009), we should not continue to use studies from limited habitats and species to represent trends across much broader areas or groups. In conclusion, more data is always better, inventory data is often (but not always) better, and functional traits can help guide assessments of data needs. Only when we acknowledge the limitations of the data in hand can we begin to fill in the gaps.

## Acknowledgements

We are grateful to all the professional and hobbyist bee collectors who contributed specimen records to the USDA NPIC museum dataset, and to the systematists and curators who identify, maintain, and database those collections. In particular, we owe thanks to Harold Ikerd and Skyler Burrows for their work on records used in this study.

J.M.M. was supported by a Graduate Research Fellowship from the University of Florida Biodiversity Institute. M.C.O. was supported by The National Science Fund for Distinguished Young Scholars (No. 31625024), and partially by the National Science Foundation of China International Young Scholars Program (31850410464) and the Chinese Academy of Sciences President’s International Fellowship Initiative (2018PB0003, 2020PB0142).

